# Thrombin activation of the factor XI dimer is a multi-staged process for each subunit

**DOI:** 10.1101/2023.02.11.528103

**Authors:** Awital Bar Barroeta, Pascal Albanese, J. Arnoud Marquart, Joost C.M. Meijers, Richard A. Scheltema

## Abstract

Factor XI (FXI), a protein in the intrinsic coagulation pathway, can be activated by two enzymes. In hemostasis, FXI is activated by thrombin, while FXIIa-mediated activation is prothrombotic. The interactions between FXI and its activating enzymes are poorly understood due to their transient nature. Here, we applied structural proteomics, molecular dynamics simulations and binding assays to investigate the interface between thrombin and FXI including the dynamics underlying FXI activation. We demonstrate that activation of FXI is a multi-staged process, where thrombin first binds to Pro520 on FXI, after which it migrates towards the activation site by engaging the apple 1 domain and finally Arg378. We validated with known mutation sites and additionally found that Pro520 is conserved in prekallikrein (PK). This enables binding of thrombin even though it cannot activate PK. Understanding the exact binding of thrombin to FXI points a way for future interventions for bleeding or thrombosis.

---

Factor XI is a protein that participates in the intrinsic pathway of coagulation. In the original model of coagulation, FXI could only be activated by factor XIIa (FXIIa) in the intrinsic coagulation pathway[1–3]. However, it was shown that FXI can also be activated by thrombin, creating a positive feedback loop that functions to maintain clot formation and stability[4–10]. A deficiency of FXI is associated with a mild bleeding disorder, while high levels of FXI are risk factors for venous and arterial thrombosis. Currently, thrombin-mediated activation of FXI is considered important for hemostasis, while FXIIa-mediated FXI activation may contribute to thrombosis.

FXI is a dimer of identical monomers, each consisting of four homologous apple domains (heavy chain) at the N-terminus that form a planar base for the chymotrypsin-like catalytic domain (light chain) at the C-terminus (Figure 1A)[11]. The catalytic domain is formed by two beta-barrels linked by a flexible loop and contains a serine-based catalytic triad. In the planar base, each apple domain is structured into an anti-parallel beta-sheet of seven beta-strands that is curved around an alpha helix[11]. The apple domains are known to mediate the interactions of FXI with other proteins or charged surfaces. Moreover, the apple 4 domain mediates the covalent assembly of a homodimer through a disulphide bridge between the Cys321 residues of the two monomers[11–13]. FXI is activated through cleavage of the Arg369-Ile370 bond in the activation loop between the apple 4 domain and the catalytic domain[11,12,14]. Initially, only one monomer is activated, generating a 1/2-FXIa intermediate that is functional and can activate FIX. The second subunit is activated in a second stage to obtain FXIa[15].

**Figure 1.**
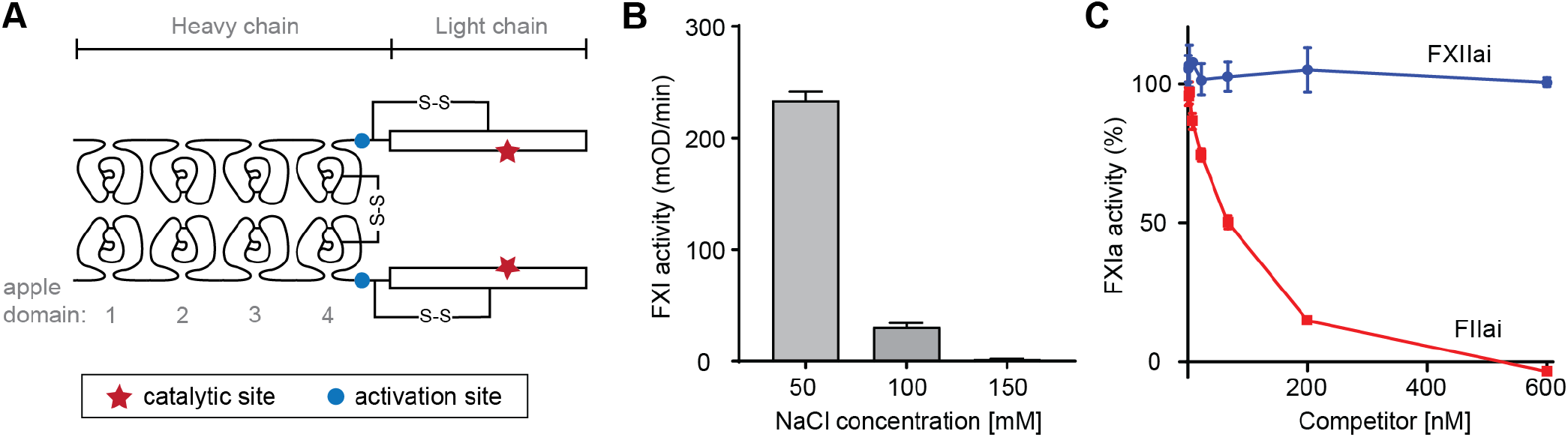
Interaction between FXI and thrombin. **(A)** Schematic model of FXI. **(B)** Salt concentrations affect FXI activation by thrombin. FXI activation was monitored in buffer containing 50 mM, 100 mM, or 150 mM NaCl. **(C)** FXI activation by thrombin in the presence of FIIai or FXIIai. Thrombin-mediated activation of FXI was monitored upon addition of increasing concentrations of inactive thrombin (FIIai) or FXIIa (FXIIai). FIIai showed a dose-dependent reduction of FXI activation, while FXIIai had no effect. Data represented mean ±SD (n=3)

The interaction between FXI and thrombin is highly transient and difficult to study. This is exacerbated by the slow conversion rate of FXI when activated by thrombin, further complicating its characterisation[4,8,10,16,17]. Nevertheless, peptide studies have proposed that thrombin interacts with the Ala45–Ser86 region in the apple 1 domain of FXI[18]. The enzymatic nature of the interaction implies a short-lived interplay between the proteins. In order to study such an interaction, it is ideally stabilized by covalent bonds between the two protein structures. Crosslinking reagents, small and agile chemicals covalently connecting amino acids in close proximity, have been developed for precisely this purpose and when combined with mass spectrometry (termed crosslinking mass spectrometry, or XL-MS) assist in the extraction of structural information for the proteins and their interactions[19]. After the crosslinking step the sample is reduced, alkylated and digested yielding 3 distinct peptide products. Of these, the crosslinked peptide pairs are structurally informative upon identification of the peptide sequences. From these sequences, the locations within the protein structures can be derived yielding information within one protein (both peptides from the same protein; intralinks) or between two proteins (peptides from different proteins; interlinks). The interlinks can be used to *e*.*g*. define the interaction interface between the two proteins and to even model the protein complex *in-silico*[20] followed by molecular dynamics (MD) to study the behaviour of the complex in solution[21]. Ultimately, the spacer length between the two reactive groups in the crosslinking reagent determines the maximum distance that can be bridged between lysine residues[22], which results in a resolution of 20 – 30 Å for this technique. Detection of the low-abundant crosslinked peptide pairs is a challenge, however, as they are present in a large background of unmodified peptides. To resolve this, enrichment handles were directly incorporated on the crosslinking reagents. One such reagent, termed PhoX, incorporates a small phosphonic acid in the spacer region that enables enrichment of crosslinked peptides using immobilised metal affinity chromatography[23]. With this efficient approach the detection of crosslinked peptide pairs becomes readily feasible as the majority of the background is removed.

In this work we report the interaction dynamics between FXI and thrombin *in vitro* and *in silico*. We find that the activation speed is sensitive to salt concentrations and is fastest at 50 mM while almost completely stagnant at 150 mM NaCl. The reduced speed of activation at high salt condition is likely advantageous for detecting the interaction with crosslinking reagents given that the crosslinking reagent takes minutes to complete. Additionally, we localize thrombin on the catalytic domain (light chain) of FXI and further use the resulting distance constraints from XL-MS to construct an initial model of the full complex. This, however, places thrombin far from the cleavage site, excluding direct activation of FXI. To investigate how this is accomplished we employed molecular dynamics (MD) simulations from which we were able to show that, after the initial binding step, thrombin engages with the previously reported apple 1 domain and two amino acids on FXI for which mutations exist that are known to lead to reduced activity. The trajectory of thrombin proposed by the MD simulations is further supported by competition studies of FXI-thrombin binding with anti-apple 1 nanobody 1C10 and prekallikrein (PK).

## Materials and methods

### FXI-WT and FXI-S557A production

FXI wild type (FXI-WT) with Cys11 replaced by Ser, and FXI-S557A were prepared as previously described [24]. The purified proteins were dialysed against 10 mM HEPES, pH 7.5, containing 0.5 M NaCl and stored at −80°C until further use. The concentration of recombinant protein was determined by measuring the absorbance at 280 nm using the extinction coefficient for FXI (13.4).

### FII-S568A production

FII-WT cDNA with Ser568 (pre prothrombin numbering, Ser525 in prothrombin numbering, Ser195 in chymotrypsin numbering) replaced by Ala was produced as previously reported [25]. Purified protein was dialysed against 10 mM HEPES, pH 7.5, containing 0.6 M NaCl and stored at −80°C. The concentration of recombinant protein was determined by measuring the absorbance at 280 nm using the extinction coefficient for FII (13.6).

### FII-S568A activation

Activation of FII-S568A was performed as previously described [25]. Purified protein was dialysed against 10 mM HEPES, pH 7.5, containing 0.15 M NaCl and stored at −80°C.

### FXII-R343A-R344A-S554A production

FXII-WT cDNA with Arg343, Arg344, and Ser568 replaced by Ala was introduced in pCDNA3.1. FXII-R343A-R344A-S554A was prepared from FXII wild type (WT) by Quikchange mutagenesis. Sequence analysis was used to confirm the introduced mutation. FXII-R343A-R344A-S554A was stably transfected in HEK293 cells using calcium precipitation. Medium containing 5% FCS (Lonza) was used to expand and grow cells in a Cell Factory (6320 cm2, Thermo Fisher Scientific) for expression. At confluence, serum-free medium (Optimem/Glutamax 1, Gibco) was added and collected every 48-72 hours. FXII-R343A-R344A-S554A was purified from the expression medium using an anti-FXII VHH column (5 mL, 15 mg VHH-F5 in-house) after concentrating the collected medium over an artificial kidney. The column was eluted with 100 mM Glycine, pH 2.7. The eluent was checked for contamination with FXIIa with an SDS gel. Purified protein was dialysed against 10 mM HEPES, pH 7.4, containing 0.15 M NaCl and stored at −80°C. The concentration of recombinant protein was determined by measuring the absorbance at 280 nm using the extinction coefficient (14) for FXII.

### FXII-R343A-R344A-S554A activation

FXII-R343A-R344A-S554A (8.6 μM) was incubated with α-kallikrein (0.43 μM, 3170AL, Enzyme Research Laboratories), PEG6000 (0.1%) and ZnCl_2_ (10 μM) for 2h at 37°C. Hereafter, α-kallikrein was removed bypassing the reaction mixture through a peptide IV column [26] and collecting the flow-through. The purified protein was stored at −80°C until further use.

### FXI activation by thrombin under varying NaCl concentrations

FXI-WT (30 nM) was incubated for 10 min at 37°C in assay buffer (30 mM HEPES, pH 7.4, 1 mg/mL BSA) containing varying concentrations of NaCl: 0.05, 0.10, or 0.15 mM. After addition of thrombin (5 nM, kind gift of the late Dr. Walter Kisiel, Albuquerque, NM, USA) the mixture was incubated for 30 min before addition of hirudin (1 μg/mL). The mixture was incubated for 5 minutes before adding S2366 (0.5 mM) and measuring absorbance for 10 min at 37 °C.

### Competition of FXI activation by thrombin with active-site mutants of thrombin or FXIIa

FXI (30 nM) was incubated with FXIIa-R343A-R344A-S554A or FIIa-S205A in assay buffer (30 mM HEPES, pH 7.4, 50 mM NaCl, 1 mg/mL BSA) for 10 min at 37°C. After addition of thrombin (1.4 nM) the mixture was incubated for an additional 10 min before addition of hirudin (1 μg/mL). The mixture was incubated for 5 minutes before adding S2366 (0.5 mM) and measuring absorbance for 10 min at 37 °C.

### PhoX concentration optimization

Freshly dissolved crosslinking reagent PhoX (50mM in anhydrous DMSO) was aliquoted in Eppendorf tubes and stored at −20 °C. FXI-S557A (5 μg, 3.46 μM, 0.3 mg/mL) and thrombin-S205A (2.3 μg, 3.45 μM, 0.13 mg/mL) were diluted in 20 mM HEPES buffer, pH 7.4, with 100 mM, 125 mM or 150 mM NaCl. Samples were incubated for 30 min at 37 °C. A PhoX aliquot was slowly heated to room temperature prior to use. The protein mixture (18 μL) was crosslinked with a concentration range of 0.25 – 2 mM PhoX over 30 min at room temperature, after which the reaction (20 μL) was quenched with 2.2 μL Tris·HCl (50 mM, pH 7.5). The crosslinked samples were subjected to SDS-PAGE (4-12% Bis-Tris gel at 200 V for 50 min in MOPS) under reducing and non-reducing conditions. The gel was stained with Imperial™ Protein stain for one hour and destained in water overnight.

### Crosslinking and sample preparation

FXI (58 μg, 3.46 μM, 0.28 mg/mL, monomer) and thrombin (26.6 μg, 3.45 μM, 0.13 mg/mL) were diluted in 20 mM HEPES, pH 7.4, with 100 mM, 125 mM, or 150 mM NaCl. Four replicates were prepared for each salt concentration, which were incubated for 30 min at 37 °C. Freshly dissolved PhoX (0.3 mM final concentration) was added and the crosslinking reaction was incubated for 30 min at room temperature. Hereafter, crosslinking was stopped by the addition of Tris·HCl (100 mM, pH 7.5) to a final concentration of 5 mM. Crosslinked proteins were denatured with urea (final concentration of 3 M) and sonicated 10 min in a bath sonicator (4 °C, on/off cycles of 30 sec). The resulting samples were simultaneously reduced with TCEP (final concentration of 10 mM) and alkylated with CAA (final concentration of 40 mM) for 1h at 37 °C. After dilution, the sample was digested by incubation with trypsin/LysC (1:25 enzyme to protein) overnight at 37 °C, and the reaction was quenched by addition of 0.5% TFA (pH *<* 2). The peptides were desalted by C_18_ using the Bravo AssayMAP platform (Agilent) and then deglycosylated and dephosphorylated overnight at 37 °C with the protein deglycosylation mix II (New England Biolabs) and Quick CIP (New England Biolabs), respectively, according to manufacturer instructions. Samples were acidified with TFA to a pH *<* 2 and desalted by C_18_ using the Bravo AssayMAP platform (Agilent) before enrichment of crosslinked peptides with Fe(III)-NTA cartridges in the Bravo AssayMAP platform as previously described[23].

### Liquid Chromatography with Mass Spectrometry and Data Analysis

The final enriched peptide mixture was resuspended in 20 μL 2% (v/v) formic acid prior to LC-MS/MS data acquisition. Mass Spectrometry data were acquired using an Ultimate 3000 RSLC nano system (Thermo Scientific) coupled to an Orbitrap Exploris 480 (Thermo Scientific). Peptides were first trapped in a pre-column (Dr. Maisch Reprosil C_18_, 3μm, 2 cm × 100 μm) prior to separation on the analytical column packed in-house (Poroshell EC-C_18_, 2.7 μm, 50 cm × 75 μm), both columns were kept at 40°C in the built-in oven for samples run on the Ultimate 3000 RSLC nano system. Trapping was performed for 10 min in solvent A (0.1% v/v formic acid in water), and the elution gradient profile was as follows: 0 – 10% solvent B (0.1% v/v formic acid in 80% v/v ACN) over 5 min, 12 - 37% solvent B over 55 min, 40-100% solvent B over 3 min, and finally 100% B for 4 min. The mass spectrometer was operated in a data-dependent mode. Full-scan MS spectra were collected in a mass range of *m/z* 350 – 1300 Th in the Orbitrap at a resolution of 60,000 at *m/z*=200 Th after accumulation to an AGC target value of 1e6 with a maximum injection time of 50 ms. In-source fragmentation was activated and set to 15 eV. The cycle time for the acquisition of MS/MS fragmentation scans was set to 3 s. Charge states accepted for MS/MS fragmentation were set to 3 - 8. Dynamic exclusion properties were set to n = 1 and to an exclusion duration of 15 s. Stepped HCD fragmentation (MS/MS) was performed with increasing normalized collision energy (19, 27, 35 %) and the mass spectra acquired in the Orbitrap at a resolution of 30,000 at *m/z*=200 Th after accumulation to an AGC target value of 1e5 with an isolation window of *m/z* = 1.4 Th and maximum injection time of 120 ms.

The acquired raw data were processed using Proteome Discoverer (version 3.0.0.400) with the integrated third-party XlinkX/PD nodes[27]. Normal peptide search was performed using the Mascot node against the full Homo Sapiens UniProtKB database (20401 entries on 29/09/2022) *in silico* digested with Trypsin/P with a minimal peptide length of six and two miss cleaved sites allowed. Cysteine carbamidomethylation was set as fixed modification. Methionine oxidation and protein N-term acetylation were set as dynamic modifications. For the search of mono-linked peptides, water-quenched (C_8_H_5_O_6_P) and Tris-quenched (C_12_H_14_O_8_PN) were set as dynamic modifications on lysine residues. Filtering at 1% false discovery rate (FDR) at the peptide level was applied. For crosslinked peptides, a database search was performed using XlinkX/PD nodes against a database the most abundant 100 proteins identified from the normal peptide search in the Mascot node, with the same protease and dynamic modifications allowed but an increased number of missed cleavages of 3. Identified crosslinks were only accepted through the Validator node at 1% FDR with a minimal score of 39 and a minimal delta score of 4. The crosslinks were mapped and visualized onto the structure using XMAS[28] within ChimeraX[29].

### Modelling of FXI dimer and docking of the thrombin binding to FXI

We used the alpha-fold 2 (AF2)[30] models for FXI, thrombin and PK in all structural analyses. To construct the homo-dimer for FXI from the available AF2 monomer, all crosslinks indicated as self-links were employed as described by Lagerwaard *et al*[28]. Evidence of mutual interactions between the light chain of FXI stem from the reproducible detection of self-link between Lys547-Lys547 and K553-K553, and overlapping peptide pairs containing Lys547 and Lys553, unambiguously defining a dimerization interface between two light chains of the FXI dimer (Figure 2D). This evidence, however, is not in agreement with the current known structure of dimeric FXI. In order to build the updated dimer model, we first docked the two light chains using HADDOCK guided by the above-mentioned self-links. The complete FXI monomer from the PDB 6I58 was then aligned with each of the light chain domains of the dimer. The complete model was then produced by accounting for the interchain disulphide bond between the two Cys321. First, the 20 AA (positions 360-380) loops connecting the light chain to the heavy chain were removed; second, a disulfide bond between the two Cys321 of each heavy chain was formed in UCSF Chimera 1.14 [31]. Lastly, the loops between positions 360-380 were remodelled with MODELLER[32], producing the final FXI dimer model (Supplementary Figure 1B) used for subsequent docking of thrombin.

**Figure 2.**
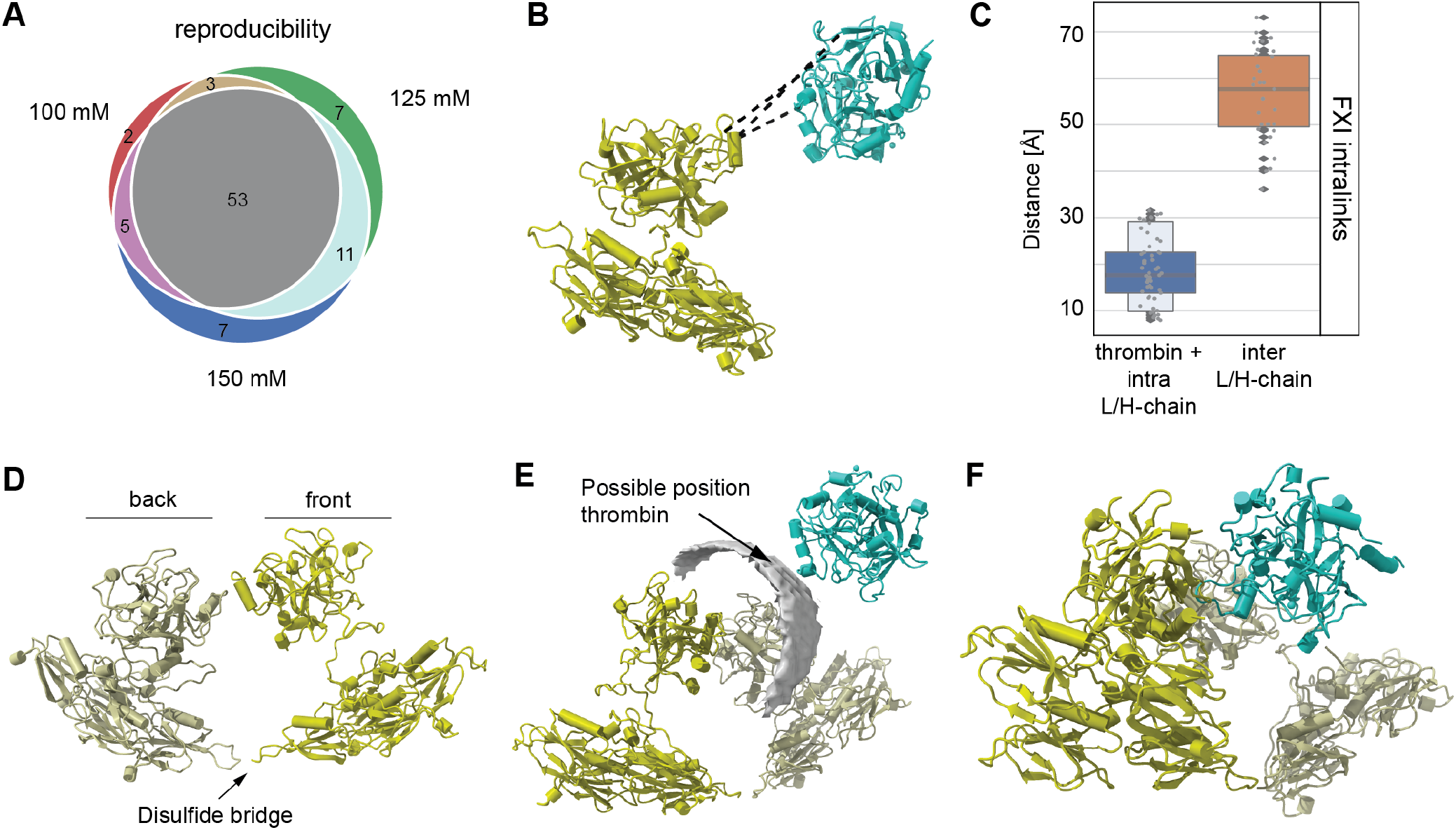
XL-MS results on FXI and thrombin. **(A)** Overlap of the detected crosslinks between the different salt concentrations. **(B)** The crosslinks (interlinks: black) mapped onto available alpha-fold 2 structures for FXI (AF-P03951-F1; forest green) and thrombin (AF-P00734-F1; sea green). **(C)** Validation of the crosslinks on FXI. **(D)** Construction of the FXI dimer (monomer #1 - forest green; monomer #2 – light green). **(E)** Possible interaction interface between thrombin and FXI (grey density). **(F)** Final model of the FXI-thrombin complex.

### Molecular dynamics simulations

The all-atom structure of the FXI dimer with docked Thrombin coarse grained (CG) was used for the MD simulations, mapping their atoms to the MARTINI 2 force field, using CHARMM-GUI online servers[33]. Starting structures contained the interchain disulfide bond between Cys339-Cys339, as well as all intra-chain disulfide bonds automatically mapped. Simulation was conducted in GROMACS 2021.4[34]. The system consisted of 3409 CG atoms for the protein representation, solvated with 42642 water beads. A neutral charge was achieved adding 468 NA (Na+) and 494 CL (Cl-), corresponding to a NaCl concentration of approximately 150 mM. The main stages of the simulation can be summarized as follows: 1) solvent and side-chain relaxation by 10,000 steps of energy minimization, imposing positional restraints of 1000 kJ mol-1 nm-2 on the whole protein. 2) solvent NVT ensemble equilibration at a temperature of 303K for 20 ns imposing positional restraints of 1000 kJ mol-1 nm-2 on the backbone beads. 3) production simulation of 180 ns in NPT ensemble at 303 K and 1 bar. Non-bonded interactions were treated with a 1.2 nm cutoff and PME for long-range electrostatics. An integration time-step of 20 fs was used during MD production runs. The system pressure was controlled by the Parrinello-Rahman barostat[35] with a coupling time of 4 ps. To analyze the trajectories, the CG structures trajectories were back mapped to atomistic detail at every 10 ns of the simulation (18 extracted snapshots in total) using the “backward” script[36].

### Generation of a nanobodies targeting the apple 1 domain of FXI

FXI and FXIa were generated as previously described [24]. Hereafter, two llamas were immunized with the coagulation factors by their subcutaneous injection. Peripheral blood lymphocytes of the immunized llamas were collected to isolate RNA and constructing a bacteriophage library as described by de Maat *et al*.[37] After an initial selection of the nanobodies on immobilised FXI, using the same procedure as described by de Maat *et al*., nanobodies were screened for FXI binding using the solid-phase binding assay described in [38].

Selected nanobodies were produced in an overnight culture of infected *E. coli* TG1 in Yeast-Tryptone medium with ampicillin (100 μg/mL) and glucose (0.2% m/v) after induction with IPTG (0.1 M final concentration). Hereafter, nanobodies were purified with a 96-wells HIS-TRAP plate or a Cobalt-NTA (Talon) column (BD Biosciences), eluting with a buffer containing 25 mM HEPES (pH 7.8), 500 mM NaCl, and 125 mM imidazole. Following a buffer change to HBS (10 mM HEPES, 150 mM NaCl, pH 7.4) using a PD10 column or a 96-wells PD G25 plate, fractions containing nanobody were combined and concentrated with an Amicon filter (MWCO 10 kDa). A NanoDrop 1000 was used to measure the OD_280nm_ and determine the final nanobody concentrations. Purified nanobodies were used to perform the following assays.

Surface plasmon resonance (SPR) experiments, performed on a BIACORE T200 (Cytiva formerly GE Healthcare) as explained in [24], were used to characterize the affinity of the nanobodies for FXI, FXIa, and individual apple domains. Following experiments were continued with nanobodies 1C10 and 2E4 that target the apple 1 domain.

### Characterization of nanobody binding epitope on FXI

The binding epitope of nanobody 1C10 on FXI was determined using hydrogen-deuterium exchange mass spectrometry (HDX MS). HDX MS was performed essentially as described in [24]. Protein samples (2 μM FXI-N72QN108Q or 2 μM FXI-N72QN108Q with 6 μM nanobody) were placed in a LEAP PAL pipetting robot (LEAP Technologies, Morrisville, NC, USA). Samples were diluted 10 times in binding buffer (10.56 mM HEPES pH 7.4, 150 mM NaCl, and 7.78 mM CaCl_2_ in 98% D_2_O), resulting in a final buffer composition of 10.5 mM HEPES pH 7.24, 150 mM NaCl, and 7 mM CaCl_2_. Samples were incubated for 10, 30, 100, or 300 s at 24 °C. Incubation with binding buffer containing H_2_O instead of D_2_O was used as reference. Deuterium exchange was quenched by mixing the sample 1:1 with quenching solution (1 g TCEP-HCl dissolved in 2 mL 2 M Urea, pH 2.5) for 1 min at 4 °C. The sample was digested by passing it over a Poroszyme Immobilized Pepsin Cartridge with an isocratic flow of 5% acetonitrile, 0.1% formic acid for 5 minutes at 4 °C. After collection on a trap (Acclaim Guard Column. 120, C_18_, 5 μm, 2.0×10 mm), the peptides were washed for 30 s at 4 °C. Subsequently, peptides were eluted and passed over a C_18_ column (Hypersil Gold C_18_: 30 mm length, 1 mm diameter, particle size 1.9 μm, Thermo cat #25002-031030) using a gradient from 0.08-64% acetonitrile at 50 μl/min at 4°C. Peptides were injected online into an LTQ Orbitrap-XL operating in positive mode. To identify peptides and their retention times, peptides were fragmented by collision induced dissociation (CID). Sequence and retention times of non-deuterated peptides were analysed with PEAKS (version 7.0, Bioinformatics Solution Inc). Deuterated peptides were analysed using HDExaminer 2.2.0 (Sierra Analytics), which calculates the deuterium uptake of peptides within 1 minute retention time. Identified peptides were assessed manually. Peptides were discarded if not accurately detected in at least half of the measurements for both compared samples. The percentage of deuterium uptake (%D) was calculated for each peptide relative to the theoretical maximal amount of deuterium incorporation. The presented HDX data consist of the mean value and corresponding standard deviation of three to five independent experiments calculated individually for four different incubation times (10, 30, 100, or 300 s). Data points with a standard deviation *>* 5% were not included.

### Functional characterization of anti-apple 1 nanobodies

Chromogenic and plasma-based assays used to characterize the effect of 1C10 and 2E4 on FXI function were performed as reported [38] unless otherwise indicated. Chromogenic assays were used to measure amidolytic FXIa activity, FIX activation by FXIa, and FXI activation by FXIIa or thrombin. Activation of FXI by FXIIa was essentially executed as described [38] but using an assay buffer composed of 30 mM Hepes, pH 7.4, 150 mM NaCl, 0.1% BSA. Plasma-based coagulation assays included activated partial thromboplastin time (APTT), factor XI activity and Calibrated Automated Thrombogram® (CAT) assays.

Activation of FXI by thrombin was monitored in a chromogenic assay as follows. FXI (30 nM) and nanobody or assay buffer (30 mM Hepes, pH 7.4, 50 mM NaCl, 0.1% BSA), were incubated for 10 min at 37°C. After addition of thrombin (5 nM) the mixture was incubated for an additional 10 min before adding hirudin (1 μg/mL). This was incubated for 5 min before adding S2366 (0.5 mM) and measuring absorbance at 405 nm for 10 minutes at 37°C.

### Fluorescence resonance energy transfer (FRET) assay

Competition of prekallikrein (PK, HPK393AL, Enzyme Research Laboratories) or nanobody 1C10 for thrombin binding to FXI was performed essentially as previously described[39] using a 20h incubation. Nanobody 2E4 was used as a negative control.

## Results

### FXI activation by thrombin as a function of NaCl

The interaction interface between FXI and thrombin is not yet well defined[4,8,10,16,17], which can be attributed to the transient nature of the enzymatic reaction. Investigation into the buffer conditions showed that FXI activation speed is dependent on the salt concentrations in the assay buffer (Figure 1B). The activation speed was significantly enhanced in assay buffer containing 50 mM NaCl compared to 100 mM or 150 mM. Moreover, this was achieved after 10 minutes of incubation, while effective activation of FXI requires more than 72 hours incubation in assay buffer with 150 mM NaCl[17]. The longer activation time observed with 100 mM or 150 mM NaCl is likely advantageous for the following crosslinking step as this takes minutes to complete.

### FXI activation by thrombin in presence of thrombin/FXIIa active site mutant

FXIIa and thrombin both activate FXI by cleaving the Arg369-Ile370 peptide bond[8,12,40], but are thought to engage different binding epitopes[18,41]. We tested this hypothesis by competing thrombin-mediated FXI activation with increasing concentrations of active-site mutants of thrombin (FIIai) and FXIIa (FXIIai). While we observed a concentration-dependent reduction in FXI activation for FIIai, the presence of FXIIai did not affect the enzymatic reaction (Figure 1C). This suggests that the enzymes do not share a binding epitope.

### Mapping of crosslinks on the FXI and thrombin protein structures

To elucidate the thrombin binding epitope we next applied crosslinking in buffers containing 100, 125 and 150 mM NaCl. These concentrations were selected to ensure that activation is still there, but at an extremely slow rate more in line with the crosslinking reaction dynamics that take minutes to complete. This compromise ensures that crosslinks between the two subunits can form and structural data can be extracted. Crosslinking was observed for all tested crosslinker concentrations without over-crosslinking (Supplementary Figure 2). We surprisingly did not observe significant differences between the crosslinking reaction using different NaCl concentrations. We did observe smearing of FXI for the lower salt concentrations, which is likely caused by improved accessibility of lysines to form mono-linked products (one reactive group of the crosslinker reacted with a lysine while the other quenched on Tris or H_2_O). To remain on the conservative side, we performed further crosslinking experiments at the lower concentration of 0.3 mM PhoX. A total of 63 (intra: 61, inter: 2) for 100 mM, 74 (intra: 72, inter: 2) for 125 mM, and 76 (intra: 74, inter: 2) for 150 mM were identified from the acquired mass spectrometry data. The different salt concentrations gave surprisingly reproducible identifications (Figure 2A). As expected, we detected the fewest links for 100 mM with increasing numbers for the higher salt concentrations. Going further, the data from all salt concentrations were used, especially as a unique interlink was detected in the 100 mM dataset bringing the total number of unique interlinks between FXI and thrombin to 3.

As quality control for the detected crosslinks, we mapped the intralinks on alpha-fold 2 structures of thrombin and FXI (Figure 2B). All links in thrombin and within the light and heavy chains of FXI fall well within the maximum distance constraint of 30 Å (Figure 2C). There is, however, a set of 30 intralinks that exceeded the maximum distance constraint. These were exclusively between the light and heavy chain of FXI and located around Lys183 and Lys248 on the heavy chain on the outer edge of the protein structure (Supplementary Figure 1A). As they are localized on the outer edge of FXI, it is very likely that these intralinks formed, because the chains can reach each other through the flexible linker region connecting them and we excluded them from further analysis. The existence of such conformations, induced by the flexible linker region, is well supported by reported electron microscopy data[42].

### Constructing the complete model

A total of 5 self-links could be extracted from the crosslinking results. Such links are intra-links (*i*.*e*. both peptides are from the same protein), but necessarily are between two distinct copies of that protein and can be defined as interlinks as described by Lagerwaard *et al*[28]. We used these links together with the disulfide bridge at Cys321 to construct the dimer using the *insilico* structural modeling tool HADDOCK[43]. The resulting model places the two subunits mirrored to each other, which is the only possibility to satisfy the disulfide bridge, the self-links and adjacently place the light chains (Figure 2D). Such a mirrored construction is likely as it automatically enables the previously reported two-stepped activation procedure[15]. The final model slightly rotates the light chain of FXI compared to the crystal structure (Supplementary Figure 1B). This is, however, well within the range of possibility for the flexible linker connecting the light and heavy chains.

To construct the final model of FXI interacting with thrombin, we used a set of computational structural modeling tools. First, we investigated the detected interlinks with DisVis[43]. Interlinks that do not agree with the majority can be excluded from this analysis, which in this case was none. Additionally, the center of mass of thrombin relative to FXI can be detected (Figure 2E; grey density). From this, it is clear that thrombin can interact with the FXI dimer on both sides of the light chain. This was also expected, given the mirror construction of the homo-dimer that enables access from either side to one of the monomers. Next, we applied protein-protein docking by HADDOCK[43] to obtain the final model using the 3 interlinks. This resulted in a model where thrombin is located on the light chain of one monomer (Figure 2F). The location is surprising however, as this places thrombin distal from the apple 1 domain on FXI, a domain that was previously reported as key for thrombin activation[18], as well as from the activation site.

### MD simulation

Our crosslinking results place thrombin on the light chain, which likely represents the initial (most stable) contact point for thrombin. Thrombin then needs to migrate towards the cleavage site to be able to activate FXI, which represents a highly dynamic process. This is less easily caught through crosslinking because this reaction takes minutes, while activation is likely occurring at nanosecond levels. To study the dynamics behind the FXI-thrombin interaction we employed MD simulations. From the results it can be observed that the structure of FXI first compresses (Figure 3A), after which thrombin migrates towards the cleavage site.

**Figure 3.**
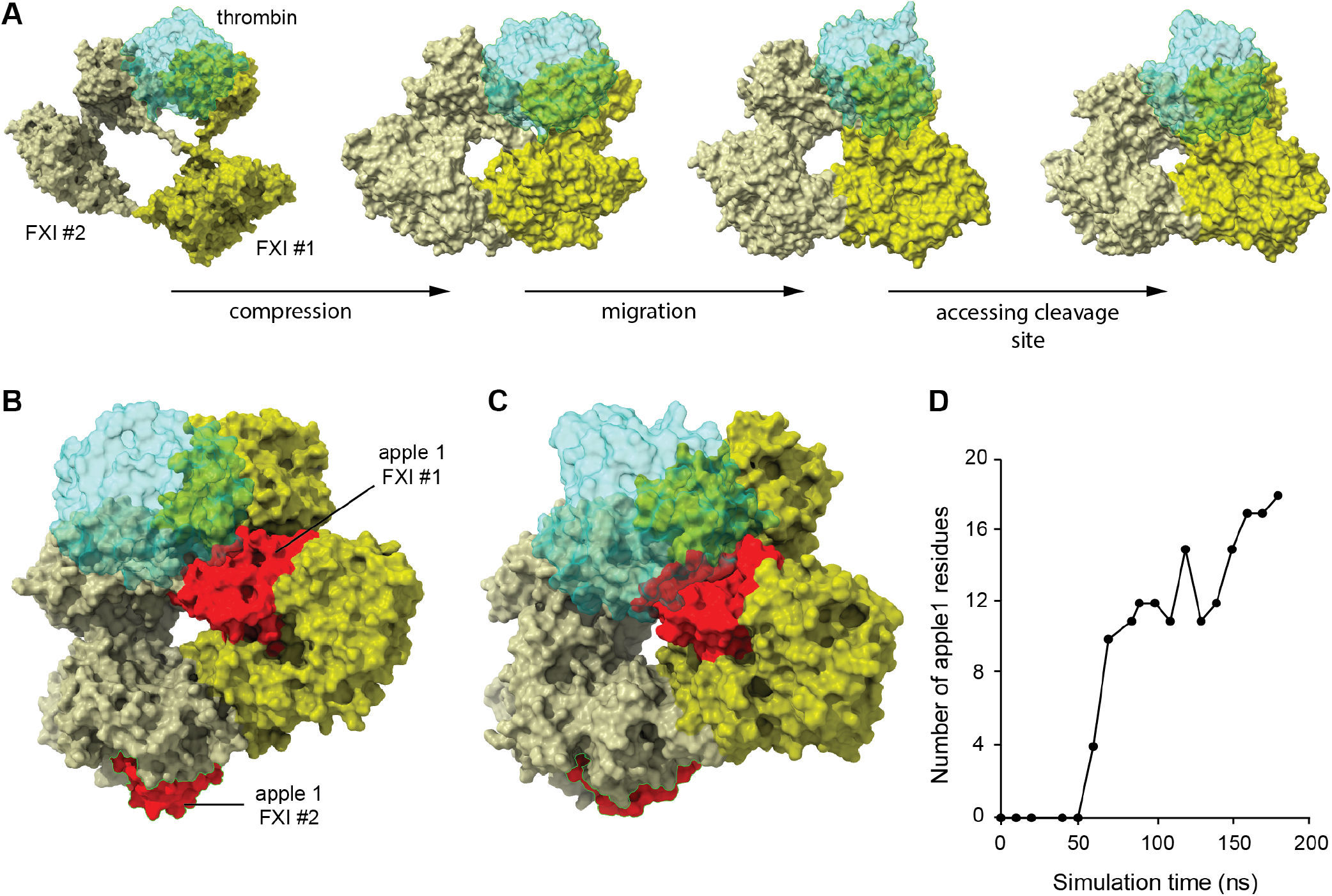
Molecular dynamics simulations. **(A)** Molecular dynamics simulation shows that upon binding the FXI/thrombin complex undergoes three stages: compression, migration, and finally cleavage site accession. **(B)** Engagement of the apple 1 domain at 60ns in the simulation or during migration. **(C)** Increasingly more protein surface of FXI connected to thrombin. **(D)** Number of residues in the apple 1 domain contacting thrombin over a simulation time of 180 ns.

At the start of this migration, thrombin engages the apple 1 domain of the FXI subunit (Figure 3B). This domain was previously reported as key for thrombin binding via amino acids Ala45–Ser86[18]. From the MD simulations amino acids Ala45–Ile76 are highlighted as contacting as well as residues Cys2–Gln5. The N-terminal residues are near to the others and potentially also play a structural role. Based on these results, we conclude apple 1 is important to initiate the migration of thrombin towards the cleavage site but not for the initial binding step. This is supported by the observation that more surface area of the apple 1 domain becomes engaged with thrombin over time (Figure 3C,D).

### Effect of naturally occurring mutations on FXI

A total of 185 mutations in FXI have previously been described[44–46]. The biggest fraction of these influence levels of FXI (n=88), and the dimerization state of FXI (n=9), which are not relevant for this study as they are unlikely to affect thrombin binding. The remaining fraction of mutations however may affect the activity or activation of FXI (n=51) (Figure 4A). To further distinguish these mutations, we verified whether the mutated sites are conserved in plasma prekallikrein (PK) (Supplementary Figure 3). This monomer homolog of FXI is not activated by thrombin [47] and therefore shared mutation sites are potentially not involved in activation by thrombin. Of the 51 combined mutation sites that affect activity, 17 are unique for FXI (Figure 4A; inset). Mapping this set of mutations on the constructed dimer of FXI shows that the majority of the unique mutations (*i*.*e*. not occurring in PK) are surface exposed (13 of 17), pointing to a possible role in binding with thrombin, while the majority of the conserved mutations are inside the globular domain, pointing to a possible role in altering the structure of FXI active site (Figure 4B). However, from the MD simulations we observed that the migration of thrombin is a highly dynamic process that exposes and hides amino acids in the process and some of the conserved sites may still play a role.

**Figure 4.**
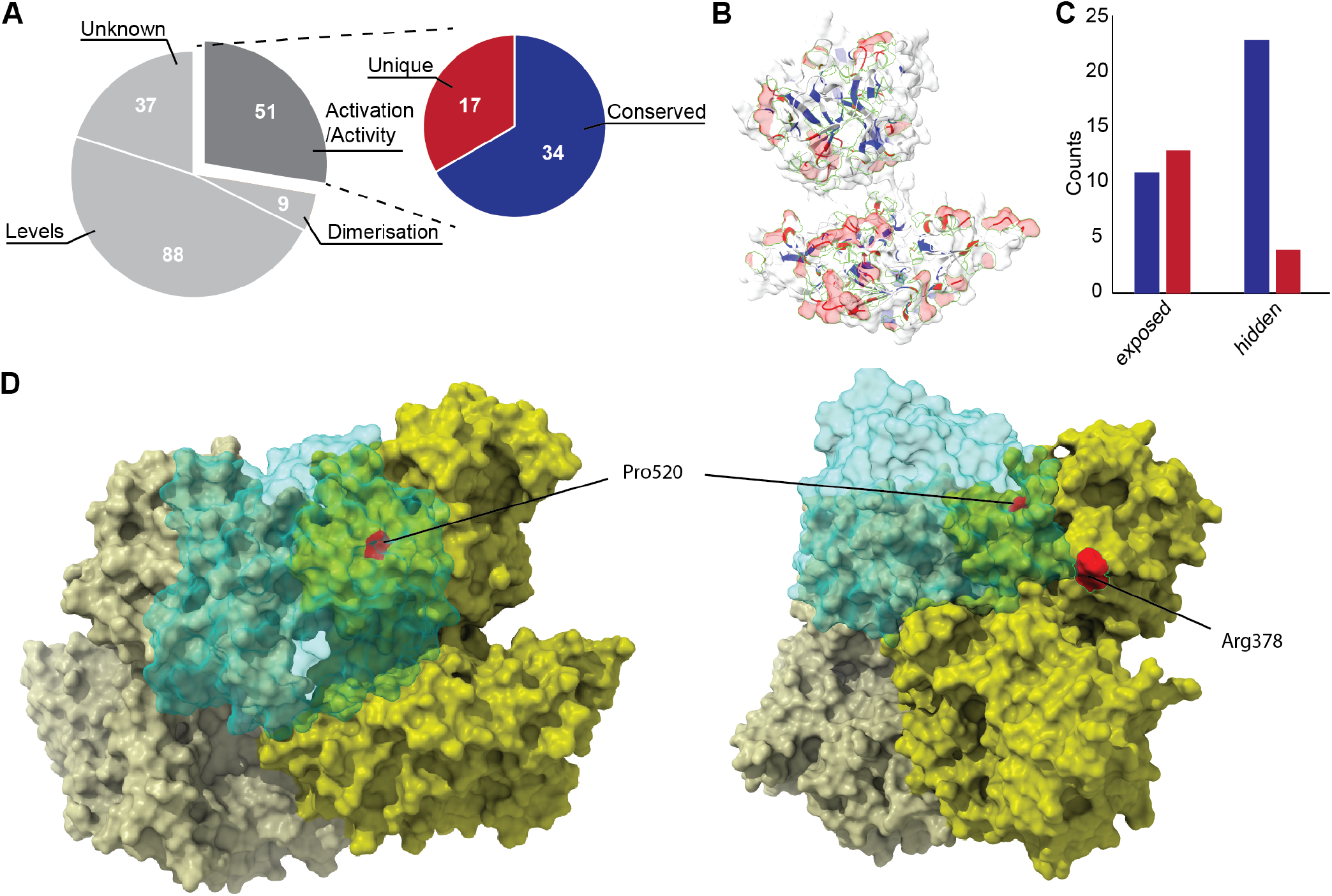
Mutations on FXI. **(A)** Effect of all known mutations. In the inset a zoom-in on the mutations known to affect the activity of FXI separated up by whether the amino acids are conserved in PK (in blue) or unique for FXI (in red). **(B)** Mutations affecting activity mapped on FXI (same color scheme as in A). **(C)** Mutations affecting activity categorized into whether they are surface exposed. **(D)** Molecular dynamics simulations uncovered Pro520 as important in binding (left; present always in the MD simulation) and Arg378 in migration (right; present after 130ns of MD simulation time)

### Competition for the FXI-thrombin interaction with nanobody 1C10

The role of the apple 1 domain in the interaction between thrombin and FXI was further investigated with nanobody 1C10. This nanobody specifically targets the apple 1 domain, as confirmed with hydrogen-deuterium exchange mass spectrometry (HDX MS). HDX MS illustrated that the binding epitope spans peptide regions Val38-Val48 and Phe61-Phe77 (Figure 5A). Although this nanobody was able to interfere with the thrombin-mediated FXI activation in a dose-dependent manner (Figure 5B), it did not affect binding between thrombin and FXI as monitored with a fluorescence resonance energy transfer (FRET)-based assay [39] (Figure 5C). This fits with the trajectory we obtained in the MD simulations, as the initial binding is far removed from the apple 1 domain and further highlights a secondary role for this domain in the interaction of thrombin with FXI that follows initial contact with the light chain. A control nanobody, 2E4, also directed against the apple 1 domain (determined by surface plasmon resonance, data not shown) did not influence thrombin-mediated FXI activation (Figure 5B) nor FXI-thrombin binding (Figure 5C).

**Figure 5.**
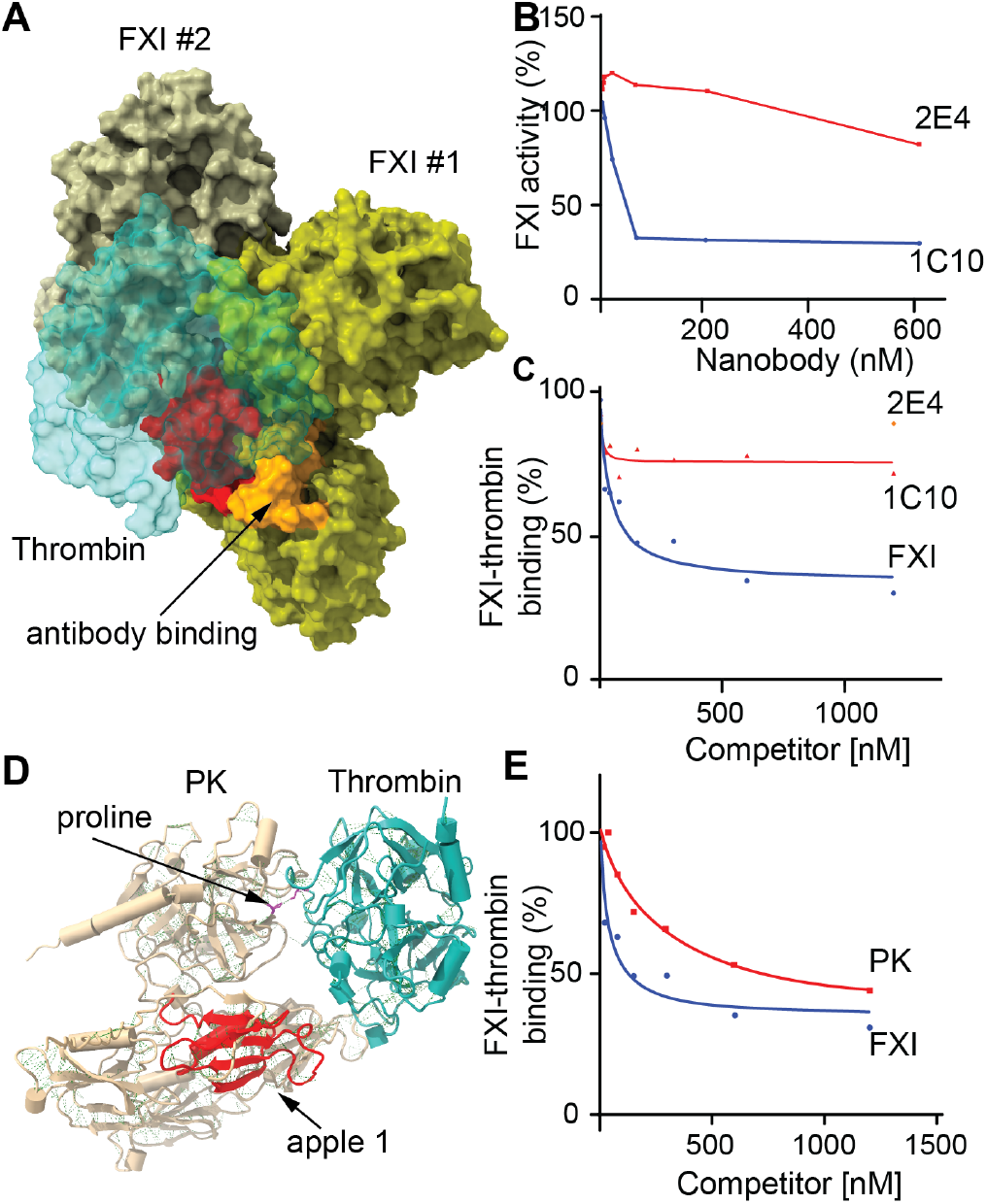
Competition of thrombin binding to FXI by PK and nanobody 1C10. **(A)** Epitope for antibody 1C10 (orange) on apple1 (red) as determined by HDX MS. **(B)** FXI activation by thrombin is dose-dependently competed by nanobody 1C10 and not by control nanobody 2E4. **(C)** Nonlabelled FXI competes, while nanobodies 1C10 and 2E4 do not, for AlexaFluor647-labelled thrombin (300 nM) binding to Europium-labelled FXI (50 pM) in a FRET assay. **(D)** Molecular docking of thrombin and PK based on the residue Pro522. **(E)** Nonlabelled PK competes with binding between FXI and thrombin in the FRET assay. Curves (B,C,E) show results from a representative experiment (n=3).

### Evolutionary conserved binding site in PK

Further investigating which FXI amino acids actually make contact with thrombin uncovered two possibly involved mutation sites. The first, Pro520, is conserved in PK but appears structurally important in binding thrombin as it remained in contact with thrombin at all timepoints of the MD simulation. This would indicate that PK may bind thrombin, even though it is not activated by it[47]. To verify whether this is possible, we performed molecular docking with HADDOCK of thrombin and PK based on the active residue Pro522 of PK. This indeed resulted in a credible model (Figure 5D). We additionally confirmed that PK can interact with thrombin by demonstrating that it competes for thrombin binding to FXI (Figure 5E) using a fluorescence resonance energy transfer (FRET)-based assay [39]. Given that the conserved proline is involved in the interaction with thrombin from the start of the simulation, binding to PK further supports that this amino acid is important for the initial binding of thrombin. Although PK also contains an apple 1 domain, it is not known whether this domain can engage in thrombin binding. The second mutation site, Arg378, is not conserved in PK pointing to a pivotal role in the actual activation.

### Subunit activation is a multi-staged process

Based on the constructed models and the MD simulations, a chain of events can be constructed that finally leads to the activation of FXI. In the initial binding step Pro520 is responsible for binding thrombin, then the apple 1 domain starts to engage to migrate thrombin towards a location between the light and heavy chain, which is followed by Arg378 locking thrombin in place between the light and heavy chains. Locked between these two, thrombin can start to activate FXI.

## Conclusions

The binding site of thrombin in FXI has remained elusive due to its short-lived nature as an enzymatic interaction. Moreover, conversion to FXIa by thrombin is very inefficient, implying a poor interaction that obstructs proper characterization[4,8,10,16,17]. Little is therefore known about the interaction site of thrombin on FXI, but it has been proposed to span Ala45–Ser86 in the apple 1 domain of FXI[18]. After binding, thrombin has to access the Arg369-Ile370 bond in the activation loop between the apple 4 domain and the catalytic domain to activate FXI[11,12,14]. Here, we report the use of crosslinking mass spectrometry and integrative modelling to elucidate the interaction of thrombin with FXI.

To aid the characterization of the FXI-thrombin interaction, we investigated conditions to influence FXI activation by thrombin. We observed that reducing the salt concentration significantly sped up the activation of FXI by thrombin. Higher salt concentrations potentially shield charged residues on the protein surface, thereby hindering electrostatic interactions between FXI and thrombin[48,49]. Although higher salt concentrations were reported to alter dimerization of the apple 4 domain and reducing dimer formation[50], dimerization did not differ greatly for the salt concentrations used in the crosslinking experiments[50]. To ensure the best likelihood of success for the XLMS approach given the relatively slow speed of the crosslinking reaction, we selected the salt concentrations at the low end of speed of activation (100, 125, and 150 mM).

From the XLMS results a total of 53 crosslinks were detected in all sample conditions with excellent overlap for the higher salt concentrations where the activation is the slowest (125 and 150 mM). From the final set of crosslinks, a total of 5 self-links could be identified that together with the known disulfide bridge on Cys321 allowed us to construct the FXI homodimer[11]. The 3 interlinks between FXI and thrombin allowed construction of the full complex. Contrary to the crystal structure of the FXI dimer, in which the catalytic domains are located on either side of a central “tipi” structure formed by the two heavy chains [11], our model suggest that the catalytic domains face each other in the same vertical plane as the Cys321 disulfide bridge (Figure 2D). This conformation appears more compatible with the constitutive interaction between FXI and high-molecular weight kininogen (HK) in plasma[51,52]. HK binds the bottom plane of the apple 2 and apple 3 domains, which would sterically hinder the proximity of the heavy chain seen in the crystal structure. HK can be accommodated more readily in the model where the heavy chains lie on the same horizontal plane. HK was however not included in either the crystallization or the crosslinking experiments and it would be interesting to see whether or how HK influences FXI conformation.

The final complex model places thrombin on the light chain (Figure 2F). As this region is far removed from the activation loop, we investigated the FXI-thrombin interaction and a possible activation mechanism using molecular dynamics. Based on this, we propose that FXI compresses after binding to thrombin, shortening the distance between the catalytic domain and the apple 1 domain. This allows interaction of thrombin to the apple 1 domain, followed by thrombin moving along the heavy chain to reach the activation loop. Interestingly, this supports cis-activation (within one monomer) rather than the previously proposed trans-activation[11]. Our experimental data validated the interaction of thrombin with the apple 1 domain[18] and proposed an initial binding site of thrombin on the light chain of FXI. While FXIIa cleaves the same activation loop, its binding site has been assigned to the apple 4 domain[41]. Accordingly, FXIIai does not compete with thrombin mediated FXI activation (Figure 1C). This underlines that the enzymes have a different binding site and thus have different modes of activating FXI. Nevertheless, the interaction between FXI and FXIIa should be elucidated to confirm this.

To investigate the probability of the proposed activation mechanism of thrombin on FXI, we compared the trajectory of thrombin on FXI with mutations known to cause reduced activity of FXI in patients[44,45]. This highlighted two residues on FXI, where Pro520 is involved with early binding and Arg378 in further migrating thrombin after it engages with the apple 1 domain. Both these residues have known mutations that were shown to affect activation. Interestingly, Pro520 is conserved in PK, suggesting that thrombin can still bind this enzyme, although it cannot activate it. Indeed, we were able to show that PK competes for binding of FXI (Figure 5E). Arg378 is unique to FXI however, and likely represents a key step in the activation dynamics of FXI. Interestingly, no mutations are present on the binding interface of thrombin on the FXI apple 1 domain that lead to reduced activity. This may point to an essential role for this interface for binding other enzymes as well. A precise understanding of the interaction of thrombin, but also FXIIa, with FXI may provide essential information to guide the development of novel antithrombotic or prohemostatic treatments.

## Acknowledgments

JCMM acknowledges that this research was supported by grant 1702 from the Landsteiner Foundation for Blood Research. RAS acknowledges that this work is part of the research programme NWO TA with project number 741.018.201, financed by the Dutch Research Council (NWO). RAS further acknowledges funding through the European Union Horizon 2020 program INFRAIA project Epic-XS (Project 823839). PA acknowledge the Dutch National Supercomputer, supported by NWO, for the computational resources (grant agreement EINF-894).

## Contributions

RAS and JCMM designed the study. ABB and JAM performed the experiments. PA performed the mass spectrometry data acquisition, data analysis and the structural modelling (protein-protein docking and molecular dynamics simulations). RAS analysed the structural data. ABB, JCMM and RAS wrote the manuscript with help from all authors.

---

**Supplementary Figure 1:**
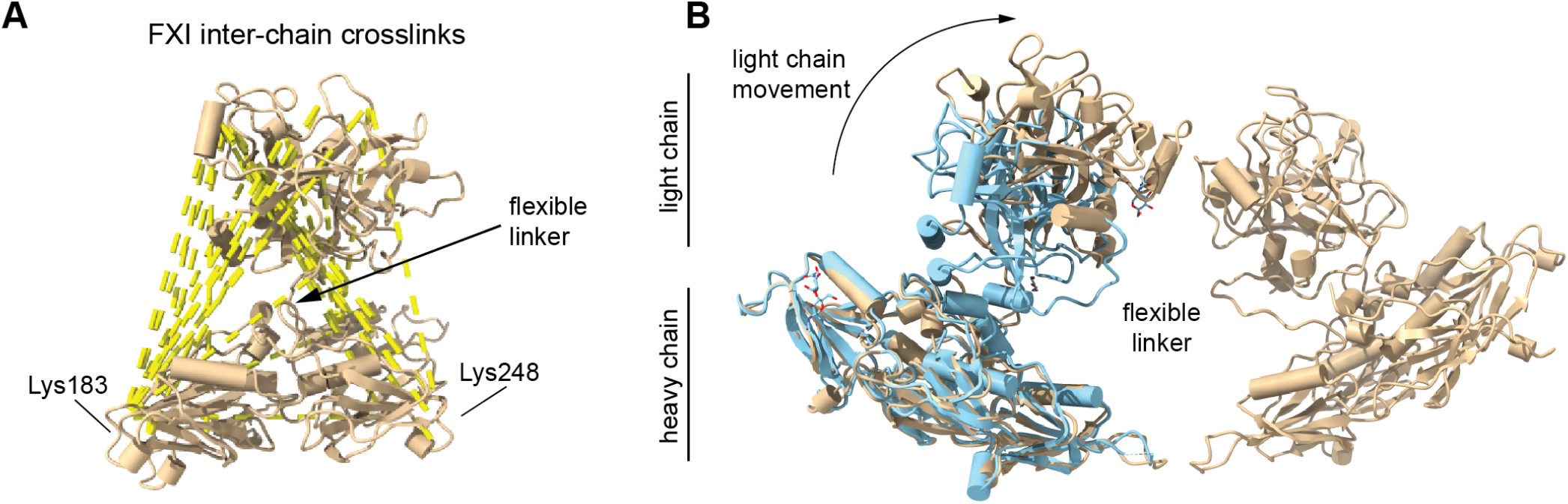
The structure of FXI. **(A)** Over-length crosslinks detected on FXI. **(B)** Overlay of the existing FXI structure (PDB: 6i58) and the constructed dimer of this work.

**Supplementary Figure 2:**
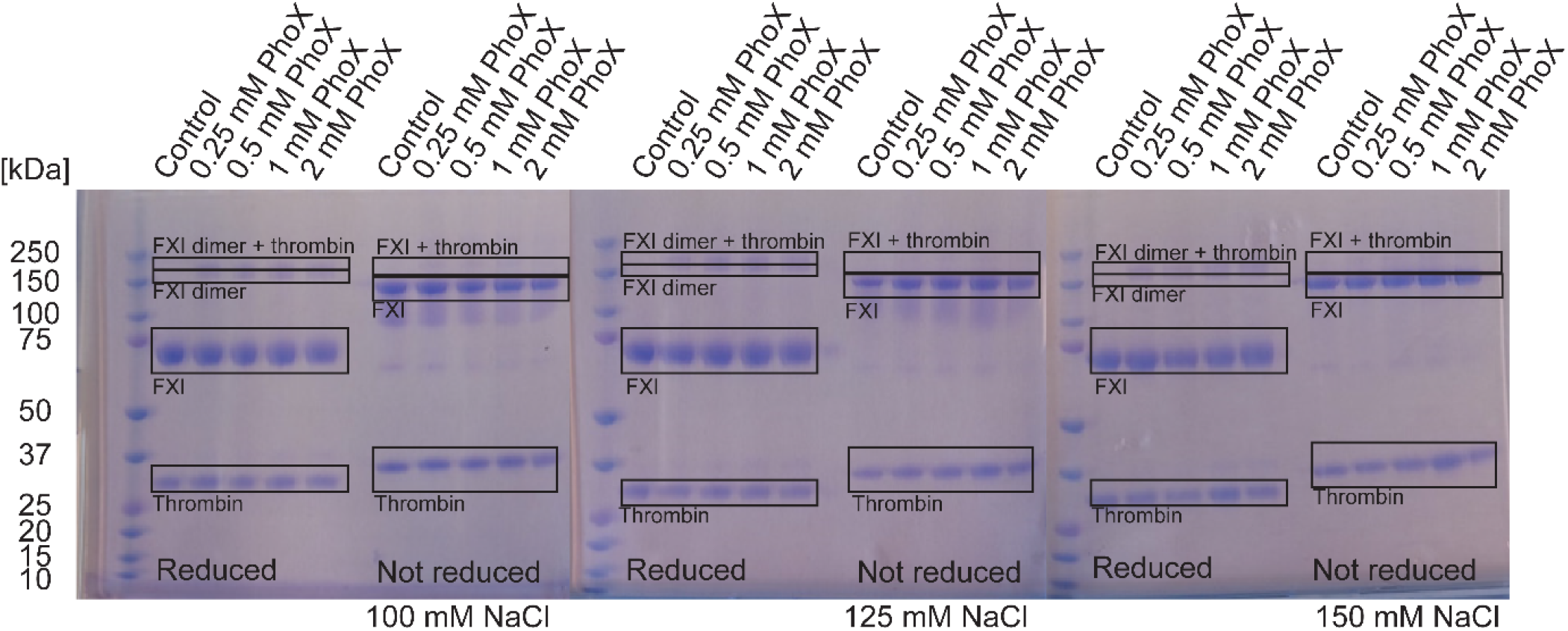
Optimisation of the crosslinker concentration for crosslinking of FXI with thrombin on SDS-PAGE. Similar read-out for crosslinking in buffer containing 100 mM (left), 125 mM (middle), or 150 mM (right) NaCl. All concentrations of PhoX show the presence of FXI dimer and FXI dimer bound to thrombin under reduced conditions. Under nonreduced conditions bands for the FXI-thrombin complex are seen in all concentrations. To prevent over-crosslinking in the larger scale experiment, 0.3 mM PhoX was used for further experiments.

**Supplementary Figure 3:**
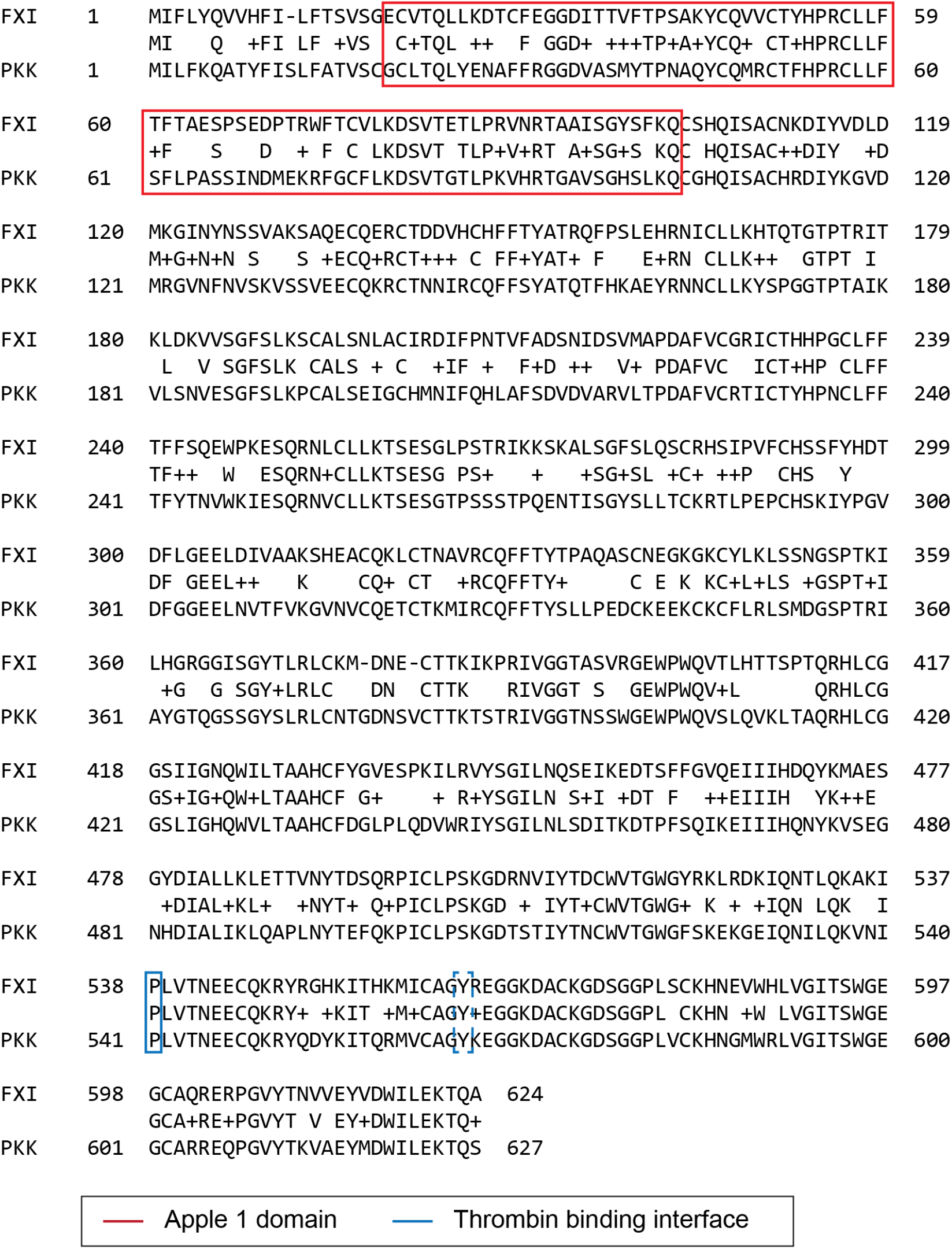
Sequence alignment of FXI and PK. The sequences exhibit distinct differences in the key apple 1 domain, but share the proline residue involved in thrombin binding uncovered from the molecular dynamics simulation.

